# Multifunctional Injectable GelMA Hydrogels with MSN-Mediated Spatiotemporal Icariin Release for Enhanced Osseointegration in Bone Defects

**DOI:** 10.1101/2025.07.16.665055

**Authors:** Yingjie Zhu, Chenyi Zhu, Yanfeng Tang, Yudong Jia, Youwen Liu, XianTao Chen, Chaowei Guo, Hongjun Li, Yuankun Zhai, Lihong Fan

**Affiliations:** Xi’an Jiaotong University, Xi’an 710004, P.R China; Medical Center of Hip, Luoyang Orthopedic-Traumatological Hospital, Luoyang, 471000, P.R China; School of stomatology HENU, Kaifeng, 475000, P.R China

**Keywords:** Mesoporous Silica Nanoparticles, Methacrylated Gelatin, Icariin, osseointegration

## Abstract

The repair of bone defects is faced with the dual challenges of limited autograft donors and single function of artificial materials. The development of new materials with mechanical adaptability, drug-controlled release and osteogenic induction has become a research hotspot in bone tissue engineering. Here, we developed an injectable photocrosslinkable hydrogel by integrating icariin (ICA)-loaded mesoporous silica nanoparticles (MSN) into gelatin methacryloyl (GelMA). The amino-functionalized MSN enhanced compressive modulus by 1.5-fold (p < 0.05) and enabled pH-responsive ICA release (91.2±4.2% cumulative release over 15 days). Our in vitro findings revealed the composite hydrogel promoted BMSC proliferation was 1.43±0.04 times that of GelMA group. And express excellent osteogenic differentiation ability (ALP activity: 3.25-fold; Mineralization: 5.01-fold). In vivo study, Micro-CT revealed significantly higher bone volume fraction (BV/TV) in rat calvarial defects at 12 weeks, with histology confirming mature trabecular bone formation. This MSN-mediated spatiotemporal delivery system synchronizes immunomodulation and osteogenesis, offering a promising strategy for non-load-bearing osseointegration.

## 1. Introduction

The high incidence of bone defects and its burden on the global public health system has become a major challenge in the field of orthopedics. More than 2 million cases of critical bone defects caused by trauma, infection or tumor resection each year [1]. However, as the gold standard therapy, traditional autologous bone transplantation still faces defects such as limited donor, secondary surgical injury and immune rejection. In addition, it also has problems such as poor shape matching [2]. Synthetic biomaterials such as hydroxyapatite and poly (lactic-coglycolic acid) (PLGA) scaffolds have been explored as substitutes. Although the shortage of donors has been partially alleviated, the lack of mechanical stability (compression modulus<5 kPa) and poor biological activity often lead to poor osseointegration [3, 4]. Therefore, the development of bone repair materials with mechanical adaptability, bioactivity and controllable degradation has become an urgent need in the field of tissue engineering.

GelMA hydrogel is a kind of gelatin derived material modified by methacrylic anhydride. In recent years, GelMA hydrogel has been widely used in bone regeneration research because of its excellent biocompatibility, tunable light polymerization characteristics and biomimetic extracellular matrix (ECM) structure [5]. GelMA is a liquid at body temperature. It is polymerized and crosslinked under ultraviolet light to form a three-dimensional porous network. The results can achieve minimally invasive injection and in situ molding, and provide customized filling for complex bone defects [6–8]. However, its inherent defects such as insufficient mechanical strength (usually less than 20 kPa) and drug release effect (such as release>60% within 48 hours) seriously limit its application in load-bearing bone repair [9]. In addition, it lacks the ability of active osteogenic induction and needs to rely on the loading of exogenous growth factors (such as BMP-2) to enhance the bone repair of induced bone defects [10]. Therefore, in order to overcome these defects, researchers try to introduce inorganic nanoparticles (such as hydroxyapatite, silica) or growth factors to enhance mechanical properties and give biological activity, but they are faced with challenges such as nano filler agglomeration, high cost of growth factors and unstable activity [11].

In order to solve the above problems, the researchers optimized the physicochemical and biological properties of GelMA hydrogel through material composite strategy. MSN has the characteristics of high specific surface area (>1000 m^2^/g), adjustable pore size (2-10 nm) and easy surface functionalization, which can be used as drug carrier and mechanical enhancement unit at the same time[12–14]. Studies have shown that MSN can enhance the mechanical properties of hydrogels through the cross-linking effect of nanoparticles. At the same time, its mesoporous channels can realize the efficient loading and continuous release of proteins, genes or small molecule drugs[15], Its ordered pore structure can also realize the slow and controlled release of drugs through pH response, such as accelerating the drug release in the slightly acidic environment of bone defects (ph∼6.5), which meets the needs of osteogenic repair in the late stage of inflammation[16]. In addition, silicic acid ions (Si^⁴+^) produced by MSN degradation can activate the expression of osteogenic related genes (such as Runx2 and Osterix) and promote the osteogenic differentiation of BMSCs[17], and bioactive silica based nanoparticles were found to stimulate the differentiation and mineralization of osteoblasts[18]. While MSNs have been incorporated into synthetic polymers (e.g.PLGA) for bone repair[19], their dispersion stability and biofunctional synergy with natural hydrogels like GelMA remain underexplored.

In order to improve the bioavailability of hydrogel, bone active substances were integrated into it. BMP-2 is an efficient growth factor, which is known to have the ability to induce stem cell differentiation and promote bone and cartilage formation. However, its high cost and potential life-threatening risk associated with its high concentration are significant disadvantages[20]. ICA as the main active ingredient of traditional Chinese medicine epimedium, has significant osteogenic and angiogenic activities. Molecular mechanism studies have shown that ICA can up regulate the expression of osteogenic markers such as alkaline phosphatase (ALP) and Osteocalcin (OCN) by activating cAMP and Wnt/β - catenin signaling pathways[21, 22]. and promote the secretion of vascular endothelial growth factor (VEGF), realizing the "osteogenic angiogenesis" coupling effect [23]. Compared with recombinant growth factor, ICA has the advantages of low cost, high stability and weak immunogenicity.

The innovation of this study is to develop a GelMA/MSN/ICA composite hydrogel system with injectability, photo responsive crosslinking and dual functionalization. The preparation process is shown in Fig 1. By covalently combining MSN loaded with ICA and GelMA gel network, a pH responsive drug repository was constructed while enhancing gel stiffness. The stepwise release of ICA (rapid release at the initial stage to inhibit inflammation and continuous release at the later stage to promote mineralization) and MSN mediated mechanical signals synergistically regulate the osteogenic differentiation of BMSCs. The purpose of this study is to systematically characterize the physical and chemical properties of the material, osteogenic differentiation effect in *vivo* and in *vitro*, and the repair efficiency of rat skull defect, verify the bone regeneration efficiency of the composite hydrogel, provide a new solution for bone defect repair, and provide a new strategy for the design of multifunctional bone repair scaffold.

**Fig. 1.**
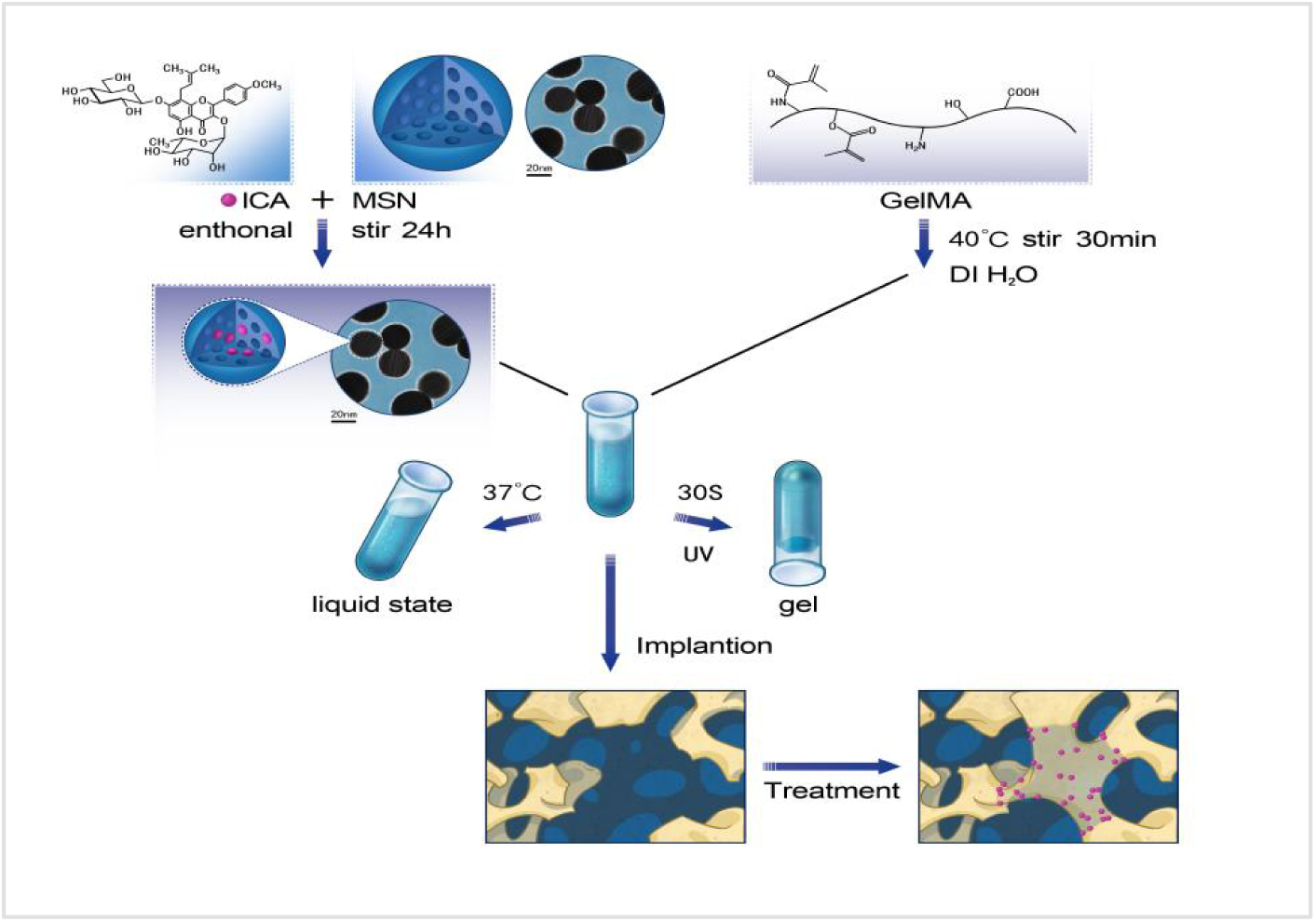
Schematic Diagram of the Composite Scaffold for Bone Defect Repair.

## 2. Materials and methods

### 2.1. ICA Loading into MSN

Amine-functionalized MSNs (100 mg) (Sigma-Aldrich) were uniformly dispersed in 0.5 mL ethanol, mixed with 5mg ICA (Sichuan, China) dissolved in DMSO, and stir at room temperature under dark conditions for 24 hours, collect drug loaded nanoparticles by centrifugation (12,000 rpm, 10 min), wash three times, and dry at room temperature. Drug loading capacity (DLC) was calculated as (mass of loaded ICA / total mass of MSN@ICA) × 100% = 2.35 ± 0.12% (*n* = 3).

### 2.2. Synthesis and characterization of GelMA/MSN/ICA hydrogels

Freeze-dried GelMA (100mg; efl-gm-60; Suzhou, China) was dissolved in DI H_2_O (1.0 ml) containing photoinitiator phenyl-2,4,6-trimethylbenzoylphosphonate (LAP) (0.3% w/v, sigma) in the dark at 50 ° C, and then the prepared MSN/ICA (0.25% w/v) was added to it, and the mixture was homogenized by ultrasonic treatment for 15 minutes (40KHz; Branson 5800) to ensure uniform dispersion. The precursor solution (200 μ L) was transferred to the test tube and crosslinked under ultraviolet light (365nm, 10m W/cm²; OmniCure S2000) for 30 seconds. The microstructure of the hydrogel was observed by field emission scanning electron microscopy (FE-SEM; Hitachi su8010). The distribution of MSN nanoparticles was determined by surface scanning using energy dispersive spectroscopy (EDS) with SEM.

### 2.3. Rheological and Swelling Characterization

The rheological properties of hydrogels were analyzed at 37 ° C using a rotational rheometer (TA instruments discovery HR-2) equipped with 20mm parallel plate geometry. The storage modulus (G′) and loss modulus (G′′) were recorded under oscillatory shear conditions (frequency 1.0 Hz, strain 1%). The gelation time is defined as G ’reaching 90% of its equilibrium modulus. The swelling rate of the composite hydrogel was measured. The pre gel solution was injected into the mold, polymerized under ultraviolet light for 30 seconds, dried and weighed (W_0_), and then the hydrogel was immersed in PBS for 24 hours and then dried and weighed (W◻) to evaluate the swelling behavior. The swelling ratio was calculated as (W◻ − W_0_)/ W_0_ × 100%.

### 2.4 Degradation rate of hydrogel

Degradation measurements were performed on days 1, 2, 3, 4, 5, 6, 8, 10, 12, 14, and 16, and residual hydrogels were weighed. Calculate the degradation rate of the hydrogel according to the weight change, and the formula is: R (%) =100 × (W_2_-W_1_)/W_1_, where W_1_ is the initial weight of the hydrogel, and W_2_ is the weight after degradation. In addition, the degradation of the hydrogel was visually assessed by rhodamine staining and assessing the degree of color fading.

### 2.5 ICA release rate

In order to evaluate the release rate of ICA from the composite hydrogel, the hydrogel was immersed in a 48 well culture plate containing 1ml Fetal bovine serum (PBS) at 37 ℃, and PBS was updated at specified intervals (1h, 5h, 12h, 1, 3, 7, 10, 14, and 16 days). Measure the absorbance of ICA in PBS at 270nm using ultraviolet (UV) spectrophotometer (lambda 800, PerkinElmer, USA). Finally, the total ICA content in composite hydrogel was quantified as 5× 10^-4^ μ mol. All experiments were independently repeated three times.

### 2.6 Cell experiments

#### 2.6.1 rBMSC isolation and culture

According to the previously described method[24]. rBMSCs were obtained from the bone marrow cavity of 3-week-old Wistar rats. rBMSCs were cultured in low-glycemic medium (Gibco, Grand Island, NY, USA) containing 10 % FBS and 1 % penicillin/streptomycin under standard culture conditions (37 °C, 5% CO_2_, and 95% relative humidity),and the culture medium was refreshed every 3 d. The cells were observed under a microscope for density and growth. If the cell density reaches around 90 %, passage at a ratio of 1:3. The third generation of rBMSCs were utilized for assessing cell viability and osteogenic differentiation. The experimental groups are: blank control group, simple GelMA group, GelMA/MSN group, GelMA/MSN/ICA group.

#### 2.6.2 Cell morphology

The morphology of rBMSCs on hydrogel was observed by cytoskeleton and nuclear staining. GelMA hydrogel, GelMA /MSN hydrogel and GelMA /MSN/ICA hydrogel were respectively placed in 48-well plate, and another group of blank control group without any material was set up, which were polymerized under UV light for 30 seconds. Then each group of samples was inoculated with rBMSCs (1 × 10^4^/well) suspension, incubated in 5% CO_2_ incubator at 37 ℃ for 24 hours, and washed twice with PBS. Then the cells were fixed with 4% paraformaldehyde (Solarbio; Beijing, China) for 30 minutes, and then permeabilized with 0.2% (V/V) Triton X-100. Then, the cytoskeleton and nucleus were stained with phalloidin and DAPI respectively according to the manufacturer’s scheme (Solarbio; Beijing, China). Fluorescence microscope (Olympus IX71, Tokyo, Japan) was used to capture the images of stained cytoskeleton (red) and nucleus (blue). In addition, after 7 days of culture, BMP-2 and Runx2 proteins in the hydrogel were stained, and the cells were stained with phalloidin and DAPI, and then observed by fluorescence microscope.

#### 2.6.3 Cell viability

The survival rate of rBMSCs in hydrogel was detected by live/dead staining (bestbio, Shanghai, China) and cell counting kit-8 (Irvine, CA, USA). GelMA hydrogel, GelMA /MSN hydrogel and GelMA /MSN/ICA hydrogel were respectively placed in 48-well plate, and another group of blank control group without any material was set up, which were polymerized under UV light for 30 seconds. Then, rBMSCs were inoculated and cultured in the corresponding 48 well plate with a density of 1 × 10^4^ cells/well for 1 and 4 days. The live/dead staining was carried out according to the manufacturer’s instructions, and observed under the fluorescence microscope. The live cells stained with calcein am emitted green fluorescence, while the dead cells stained with PI emitted red fluorescence. Image J software was used to analyze the live/dead staining images. The cell viability was further evaluated. After rBMSCs were cultured with hydrogel for 1, 3 and 5 days, CCK-8 was measured at the above time points. The absorbance was measured at 450nm using bio rad microplate reader. The cell survival rate (%) = (test od blank OD)/(control od blank OD) × 100%. All experiments were performed three times.

#### 2.6.4 Osteogenic differentiation of rBMSCs in *vitro*

The blank group, GelMA, GelMA /MSN and GelMA /MSN/ICA hydrogel were injected into 48 well plates, respectively, and polymerized under UV light for 30s. RBMSCs were seeded in the well plates at a density of 5 × 10^4^ cells/well, respectively, and cultured for 7 and 14 days. L-DMEM osteogenic medium (Cyagen, Santa Clara, CA, USA) was used to induce cell differentiation. According to the manufacturer’s instructions, BCIP/NBT staining (Beyotime, Shanghai, China) was used to observe the expression of alkaline phosphatase (ALP, Beyotime, Shanghai, China) on the 7th day, and ALP activity was evaluated with ALP detection kit (Beyotime, Shanghai, China). Alizarin red staining (ARS, Beyotime, Shanghai, China) was used to evaluate calcium deposition on day 14. After washing twice with PBS at the corresponding time, the cells were fixed with 4% paraformaldehyde (Solarbio; Beijing, China) for 30 minutes. Incubate with dye at room temperature for 30 minutes, and then wash twice with PBS to eliminate background interference. The staining and number of calcium nodules were observed under the light microscope to evaluate the osteogenic effect. The ARS stained samples were decolorized by dissolving in 10% cetylpyridinium chloride, and the absorbance at 562nm was measured by UV spectrophotometer, and the calcium deposition in each group was detected by microscope.

#### 2.6.5 Reverse transcription quantitative polymerase chain reaction (RT-qPCR)

In order to evaluate the osteogenic differentiation ability induced by composite hydrogel, the expression levels of osteogenic related genes such as alkaline phosphate (ALP), runt related transcription factor 2 (Runx2), collagen 1 (col1) were detected by RT -qPCR. The blank group, GelMA, GelMA /MSN and GelMA /MSN/ICA hydrogel were injected into 48 well plates containing osteogenic medium, respectively, and polymerized under UV light for 30s. RBMSCs were seeded into the corresponding 48 well plates (1 × 10^5^/well) and cultured for 7 days. Total RNA was extracted from rBMSCs (Invitrogen, Carlsbad, CA, USA) using Trizol reagent, and then reverse transcribed into cDNA using primescript RT Kit (Takara, Japan). RT-qPCR with Q SYBR green Supermix (Bio-Rad, Hercules, CA, USA) and a QuantStudioTM 7 Flex real-time PCR system (Applied Biosystems, Carlsbad, CA, USA). were employed to further evaluate mRNA expression levels. Relative expression levels of mRNAs were normalized to that of GAPDH and calculated by the 2^−ΔΔCt^ method. Data analysis was conducted using GraphPad software (GraphPad, Inc., La Jolla, CA, USA). All experiments were conducted three times independently. Primer sequences were designed using Primer Premier software (Premier Biosoft, Palo Alto, CA, USA), as summarized in Table 1.

**Table 1.**
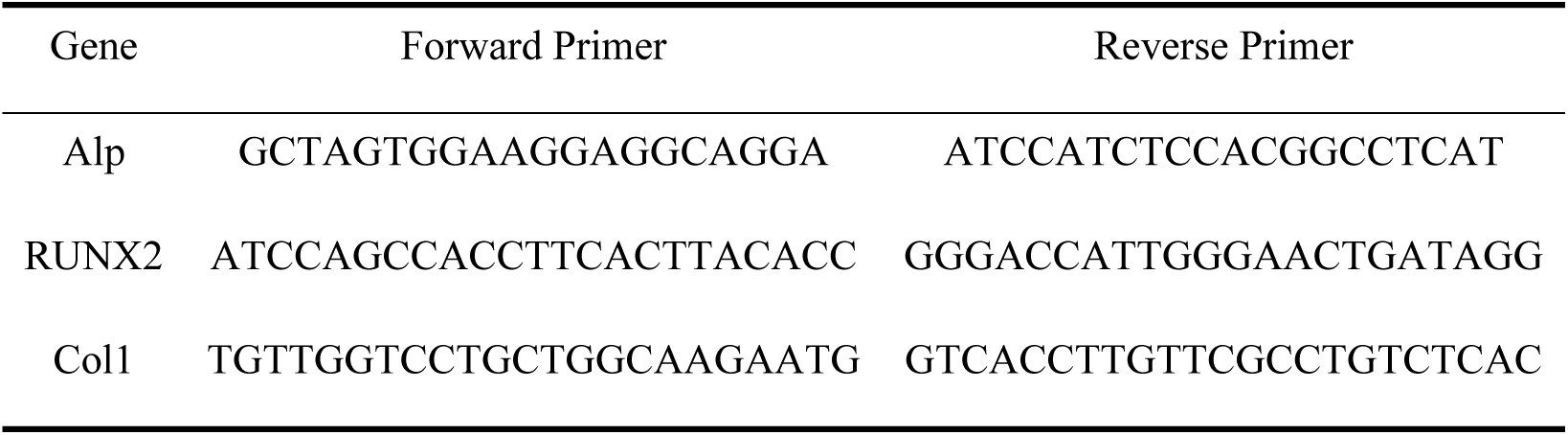
Primers used in RT-qPCR.

### 2.7 Animal experiments

All procedures involving animals were conducted in strict accordance with the ethical guidelines of Henan University School of Stomatology and approved by the animal ethics committee of Henan University School of Stomatology. A total of 40 clean Wistar rats were selected in this study. They were randomly assigned to four groups (n = 10/group) using a computer-generated randomization table", as follows: 1) blank control group (blank); 2) GelMA group; 3) GelMA/MSN group; 4) GelMA/MSN/ICA group. The rats were anesthetized by intraperitoneal injection of 3% Pentobarbital Sodium, and the rat skull defect model was prepared after anesthesia, after anesthesia, the outer side of the skull was shaved, the skin was cut to expose the surface of the skull, a 5mm*5mm bone defect of the same specification was made with a 5mm diameter ring drill, and the end periosteum was removed to prevent ossification. After repeated flushing with sterile normal saline, the hydrogel solution was dropped respectively, and the skin was sutured after 30s gelation under ultraviolet light to complete the establishment of the model. To prevent infection, penicillin (1.5 mg/kg) was administered three days a day. At 6 and 12 weeks after implantation, the rats were killed by spinal dislocation, and the bone defect tissues were collected and fixed in 4% paraformaldehyde for further analysis.

#### 2.7.1 Micro-CT analysis

In order to evaluate the bone regeneration at the bone defect site, we used the micro computed tomography (Micro**-**CT) system (Skyscan 1076 scanner, kontich, Belgium) to collect images, and used ctvox to reconstruct three-dimensional (3D) images. The captured images were reconstructed using multimodal 3D visualization software. Through image analysis, we determined several key parameters: bone volume fraction (BV/TV), trabecular thickness (TB. Th), trabecular number (TB. N) and trabecular separation (TB. SP). After Micro**-**CT scanning, the samples were decalcified in ethylenediamine tetraacetic acid (EDTA) for 8 weeks. Subsequently, the samples were dehydrated with ethanol in a graded series and embedded in paraffin.

#### 2.7.2 Hematoxylin-eosin (HE) staining

After decalcification and paraffin embedding of bone tissue specimens in each group, specimens were randomly selected from each group. The specimens were dewaxed in xylene after slicing, and then rehydrated with graded ethanol (100%, 95%, 80%, 70%, each for 5 minutes), and rinsed in distilled water. The slides were immersed in mayer’s hematoxylin (Sigma, MH532) for 8 minutes, then rinsed for 5 minutes, and differentiated in 1% acid ethanol (1% Hcl in 70% ethanol) for 30 seconds for nuclear staining. Sections were re stained with Eosin Y (0.5% 95% ethanol solution, Sigma) for 2 minutes for cytoplasmic staining. The stained sections were observed under a bright field microscope (Nikon Eclipse Ni-E) equipped with a 20x objective lens (NA 0.75) and a DS-Fi3 camera. Images were analyzed using NIS elements ar 5.21 software to assesstissue morphology, inflammation, and new bone formation.

#### 2.7.3 Immunofluorescence staining of type 1 collagen (Col1)

As mentioned above, samples were randomly selected from each group, the specimen sections were dewaxed, and the thermally induced epitopes were retrieved in a 95 ° C citrate buffer (10 mm, pH 6.0),Sigmausing Biocare medical for 20 minutes. The slides were cooled to room temperature, washed with PBS (3 × 5 minutes) and incubated with 5% normal goat serum (Vector Laboratories) in PBS containing 0.3% Triton X-100 (PBST) for 1 hour at room temperature. Col1 primary antibody (ab21286, Abcam, 1:200) diluted in 1%BSA/PBST was placed overnight in a humidifying chamber at 4 ° C. After that, the slides were washed with PBST (3×5 minutes), and incubated with Alexa fluor 488 coupled secondary antibody (Invitrogen, 1:500) at room temperature for 2 hours in the dark. The nuclei were re stained with DAPI (1μ g/ml, Invitrogen D1306) for 5 minutes, and then washed three times with PBS (5 minutes each time). The sections were covered with ProLong Gold anti-counterfeiting patches (Invitrogen) and imaged with a confocal microscope (Zeiss LSM 880 Airyscan) to observe the fluorescence intensity of Col1.

### 2.8 Statistical analysis

The sample size of each statistical analysis is n ≥ 3. All data are expressed as mean standard deviation. One way ANOVA and t-test were used to evaluate the differences between the experimental groups. SPSS software version 26.0 (SPSS Inc., Chicago, USA) was used for statistical analysis, and the significance was set to P<0.05.

## 3. Results

### 3.1 Microstructural Characterization of the Composite Hydrogel

As shown in Fig 2A, GelMA/MSN/ICA hydrogel was observed in *vitro* after preparation. It was found that the composite hydrogel was injectable at body temperature and turned to gel state after UV photocrosslinking. According to the analysis of rheological test results, as shown in Fig 2B, G’’ was higher than G’ at the beginning, indicating that the composite hydrogel was in the sol state. Under UV exposure, G’ increased rapidly, and there was a crossover point between G’ and G’’, indicating that a sol-gel phase transition occurred. As shown in Fig 2C, scanning electron microscopy (SEM) shows that GelMA/MSN/ICA hydrogel has an interconnected porous network with pore sizes ranging from 5 μ m to 20 μ M. MSN are evenly dispersed in GelMA matrix. EDS element mapping (Fig 2D) shows the presence of C, N, O and Si, which confirms the uniform distribution of collagen and Si ions on the whole composite hydrogel. At the same time, after mechanical compression, the irregular pore geometry and solid structural integrity are also maintained, which indicates that MSN integration enhances the stability. The compression modulus of composite hydrogel (12.3 ± 1.2 kPa) is higher than that of pure GelMA group (8.1 ± 0.9 kPa, ** P<0.05) was 1.5 times higher and could promote cell adhesion, indicating that MSN integration enhanced stability.

**Fig. 2.**
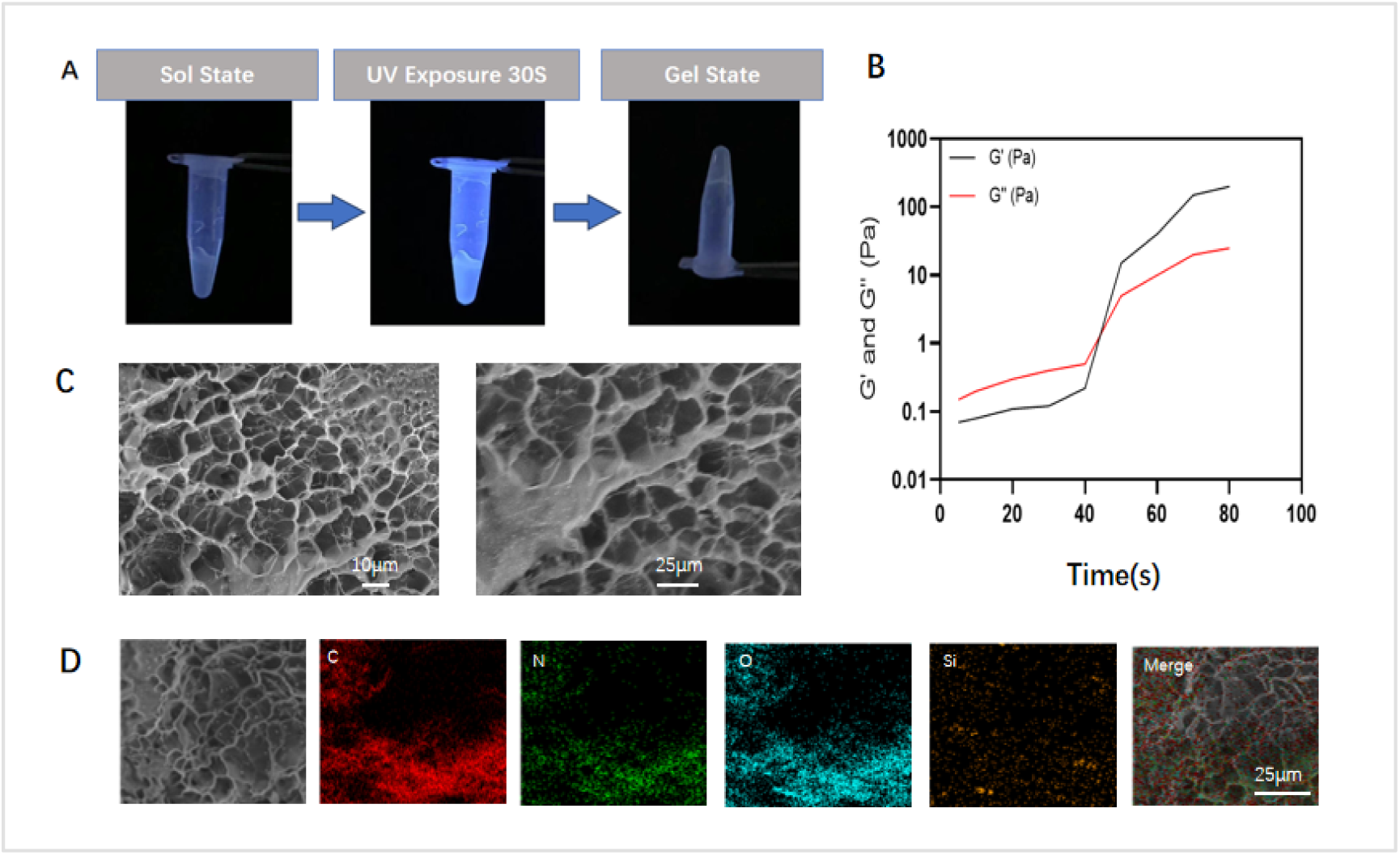
(A) Appearance of the GelMA/MSN/ICA Hydrogel, Composite hydrogel is sol state at body temperature and turned to gel state after UV photocrosslinking. (B) Rheological testing of GelMA/MSN/ICA Hydrogel. (C) Microstructure of GelMA/MSN/ICA Hydrogel. (D) EDS-SEM of GelMA/MSN/ICA Hydrogel. All values are expressed as mean ± SD, n = 3.

### 3.2 Swelling and Degradation Behavior of the Composite Hydrogel

After soaking in PBS for 24 hours, the equilibrium swelling rate of the composite hydrogel (336.8±9.6%) was significantly higher than that of GelMA hydrogel (220.7±5.6%) (Fig 3A). The addition of MSN may protect GelMA network from excessive hydration and balance hydrophilicity and structural protection. This enhanced swelling capacity indicates that the potential of nutrient exchange has been improved in a dynamic physiological environment. In *vitro* degradation of GelMA/MSN/ICA hydrogel showed that GelMA /MSN/ICA hydrogel continued to degrade within 16 days, and the degradation rate reached 94.5 ± 4.5 on the 16th day (Fig 3B), indicating that the hydrogel had good degradability. MSN component may stabilize the enzymatic hydrolysis of GelMA by collagenase, and continuously regulate the degradation of hydrogel in the process of bone repair.

**Fig. 3.**
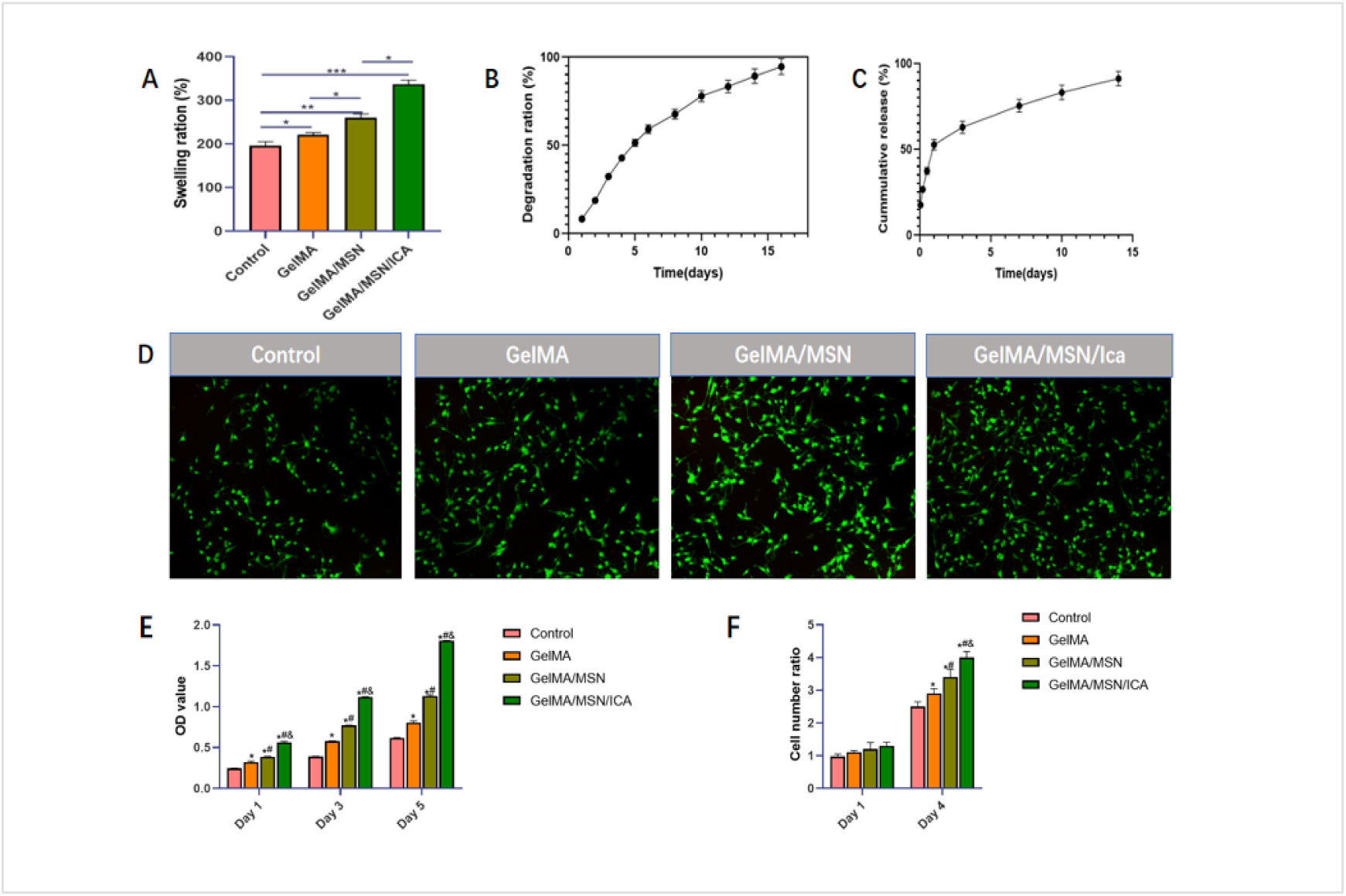
(A)Swelling ratio of GelMA/MSN/ICA Hydrogel. (B) Degradation Rate of GelMA/MSN/ICA Hydrogel. (C) Release Rate of ICA in the Composite System. (D)Live/Dead staining of rBMSCs on 4 d (Living cells were green and dead cells were red, bar = 1 mm). (E) Proliferation of rBMSCs on 1, 3 and 5d evaluated using CCK-8. (F) Cell Number Ratio of BMSCs Cultured with the control group, GelMA, GelMA\/MSN and GelMA /MSN/ICA group for 1 and 4 Days, respectively.All values are expressed as mean ± SD, n = 3.

### 3.3 ICA release rate

The encapsulated ICA was released from the hydrogel within 15 days (Fig 3C). The highest release rate was 52.7 ± 3.0% on the first day. After the initial surge, the release rate of ICA gradually decreased and reached a stable rate, which remained stable for the remaining 15 days. By the 15th day, the cumulative release rate of ICA reached 91.2±4.2%, which demonstrated the ability of MSN mesoporous channels to capture ICA molecules and regulate the diffusion, effectively prolonging the duration of ICA release, which was the key feature of long-term osteogenesis. The pH/enzyme reaction was released, which was consistent with the time requirement of bone regeneration (from inflammation stage to remodeling stage).

### 3.4 Cell morphology assessment

The effect of composite hydrogel on the morphology of rBMSCs was observed. The cells cultured in blank group, GelMA, GelMA/MSN and GelMA/MSN/ICA were stained with Phalloidin/DAPI staining (Fig 4A). In all groups, BMSCs showed complete cell morphology, and phalloidin staining revealed clear and definite cytofilament structure in each group of rBMSCs. As shown in Fig 4B and Fig 4C, through the protein staining of Runx2 and BMP-2 for each group of hydrogels, the protein staining was green. It can be seen that the expression of GelMA/MSN/ICA histone protein was higher than that of other groups, and the cell morphology was complete, indicating that the composite hydrogel would not have a negative impact on the early adhesion of rBMSCs, and has good protein expression.

**Fig. 4.**
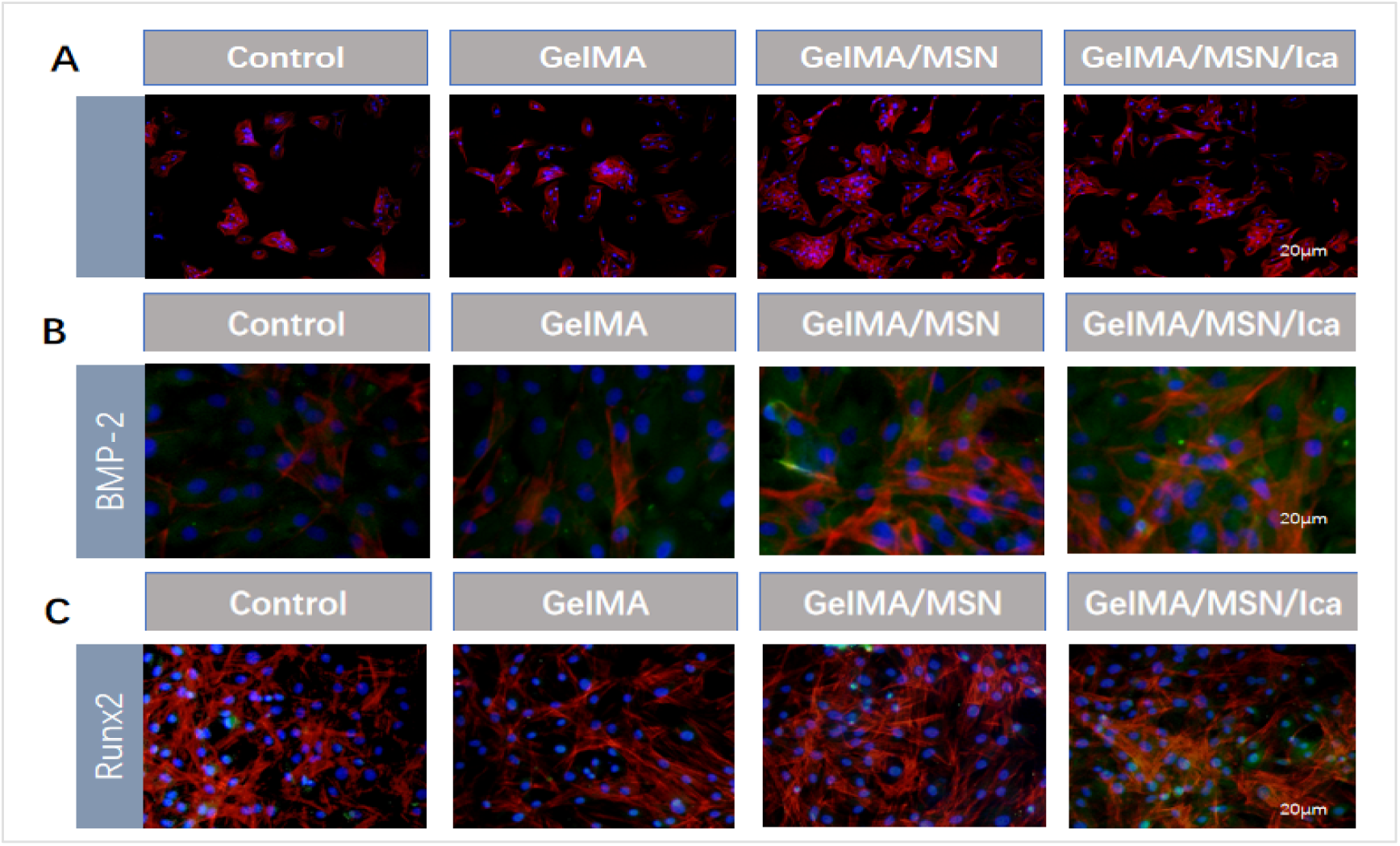
(A)Morphology of rBMSCs on 1d (Cytoskeleton was stained with Phalloidin(red) and nuclei was stained with DAPI (blue)). (B) Morphology of rBMSCs on 7d, and the protein staining of BMP-2(Green). (C)Morphology of rBMSCs on 7d, and the protein staining of Runx2 (Green).

### 3.5 Cell viability assessment

The changes of cell viability were evaluated by CCK-8 test and Live/Dead staining. CCK-8 test showed that on the fifth day, the OD value of GelMA/MSN/ICA group was 1.43±0.04 times that of GelMA group (Fig 3E). The cell density of GelMA/MSN/ICA group was the highest on the fifth day, which was significantly better than that of GelMA group, and gradually filled the whole field of vision (Fig 3D), indicating that GelMA/MSN/ICA effectively promoted cell proliferation (Fig 3F). The nano morphology of MSNs and the mitogenic effect of ICA may promote cell expansion in synergy. These findings were confirmed by Live/Dead staining, showing uniform green fluorescence (Calcein AM) and minimal red signal.

### 3.6 Osteogenic differentiation in *vitro*

Osteoblasts are derived from the differentiation of BMSCs and are the main functional cells in the process of bone regeneration. In order to evaluate the osteogenic differentiation, rBMSCs cultured in hydrogel for 7 and 14 days were detected by alkaline phosphatase (ALP) and alizarin red (ARS) staining (Fig 5A). ALP, as an early osteogenic marker, showed significant staining results on day 7 in GelMA/MSN/ICA group. After further quantitative evaluation and analysis, it was found that the alkaline phosphatase (ALP) activity in GelMA/MSN/ICA group was 1.41±0.05 times higher than that in GelMA/MSN group and 2.92±0.15 times higher GelMA group respectively (Fig 5B). On the 14th day, ARS staining showed extensive calcium nodule formation in GelMA/MSN/ICA group (Fig 5C), and early differentiation markers were mineralized before ECM.

**Fig. 5.**
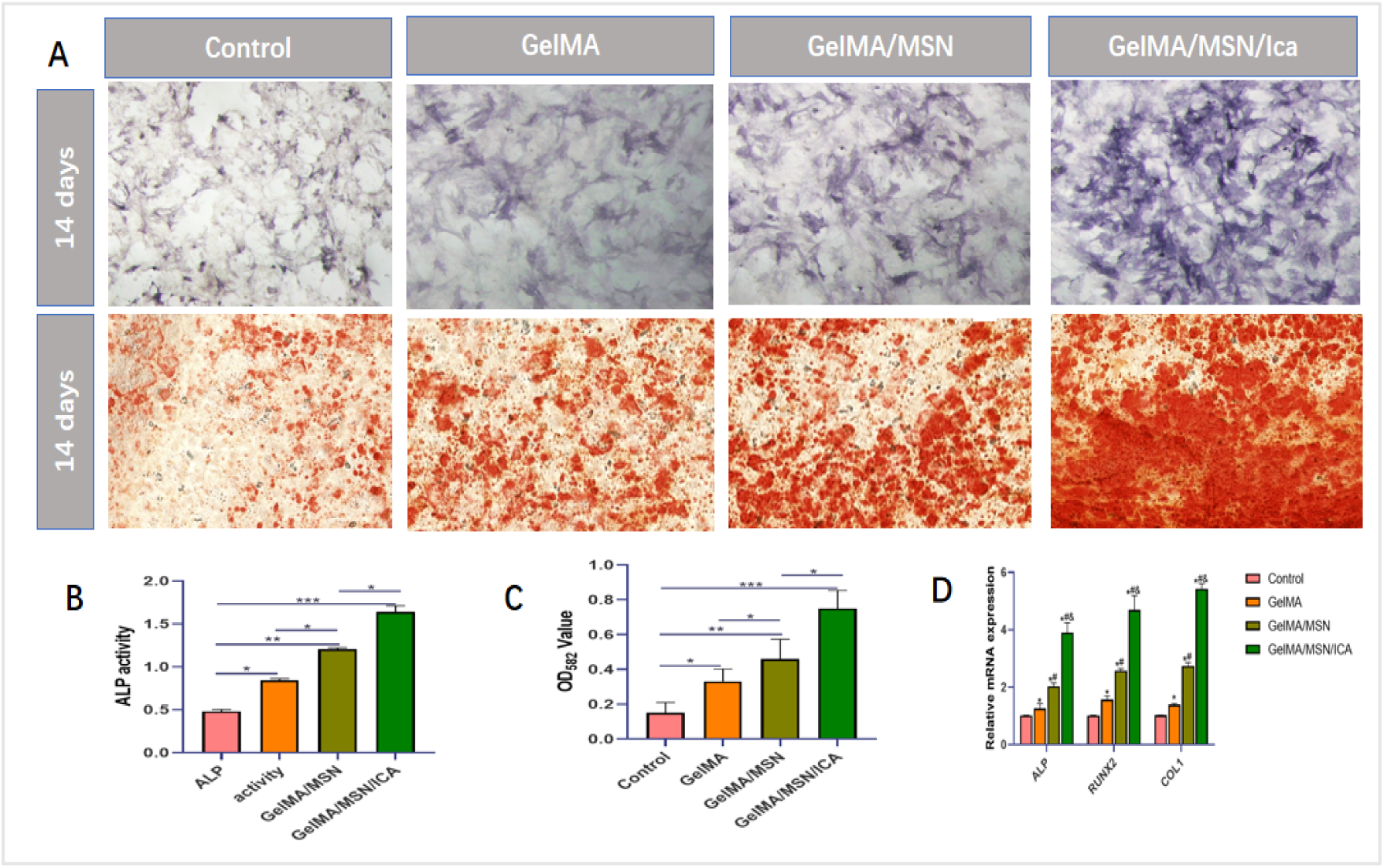
(A)Evaluation of rBMSC Differentiation Using Alkaline Phosphatase Staining and Alizarin Red Staining. (B) ALP activity of rBMSCs on 7 d. (C) Quantification of ECM mineralization on 14 d. (D) Relative fold change of osteogenic gene (ALP, Runx2, Col1) expression measured by RT-qPCR on 7 d. All values are expressed as mean ± SD, n = 3.

### 3.7 RT-qPCR analysis

RT qPCR was used to detect the expression of genes related to osteogenic differentiation in rBMSCs. The expression levels of ALP, Runx2 and Col1 genes were evaluated to clarify the effect of composite hydrogel on osteogenic differentiation of rBMSCs. According to the results, on the 7th day, the expression of ALP gene in GelMA/MSN/ICA group was 1.91±0.04 times higher than that in GelMA/MSN group and 2.87±0.08 times higher than that in GelMA group, respectively. Runx2 and Col1were also highly expressed in the composite hydrogel group as in other groups (Fig 5D). In conclusion, RT-qPCR results showed that GelMA/MSN/ICA significantly enhanced the osteogenic differentiation of rBMSCs. These results may be related to the sustained release of ICA and the synergistic activation of BMP-2/Smad and Wnt/β-Catenin pathways by MSN derived silicic acid.

### 3.8 Animal experimental evaluation

During 6 weeks to 12 weeks after operation, no infection or death occurred in the whole observation cycle. Bone tissues with the same size range including bone defects were randomly collected from each group for analysis. We observed and analyzed the bone ingrowth at the skull bone defect through Micro-CT images, as shown in Fig 6A. At week 6, bone regeneration in GelMA/MSN/ICA group was slightly higher than that in GelMA/MSN group, and significantly higher than that in GelMA group. At the 12th week, continuous bone ingrowth was observed in each group, but the bone regeneration in GelMA/MSN/ICA group was better than that in other groups, and the difference between groups was expanded. In addition, quantitative display by Micro-CT (Fig 6B) showed that the bone volume fraction (BV/TV), trabecular thickness (Tb.Th) and trabecular number (Tb.N) and other parameters in the composite hydrogel group at 6 and 12 weeks were significantly higher than those in GelMA group and GelMA/MSN group, and the gap between groups gradually increased.

**Fig. 6.**
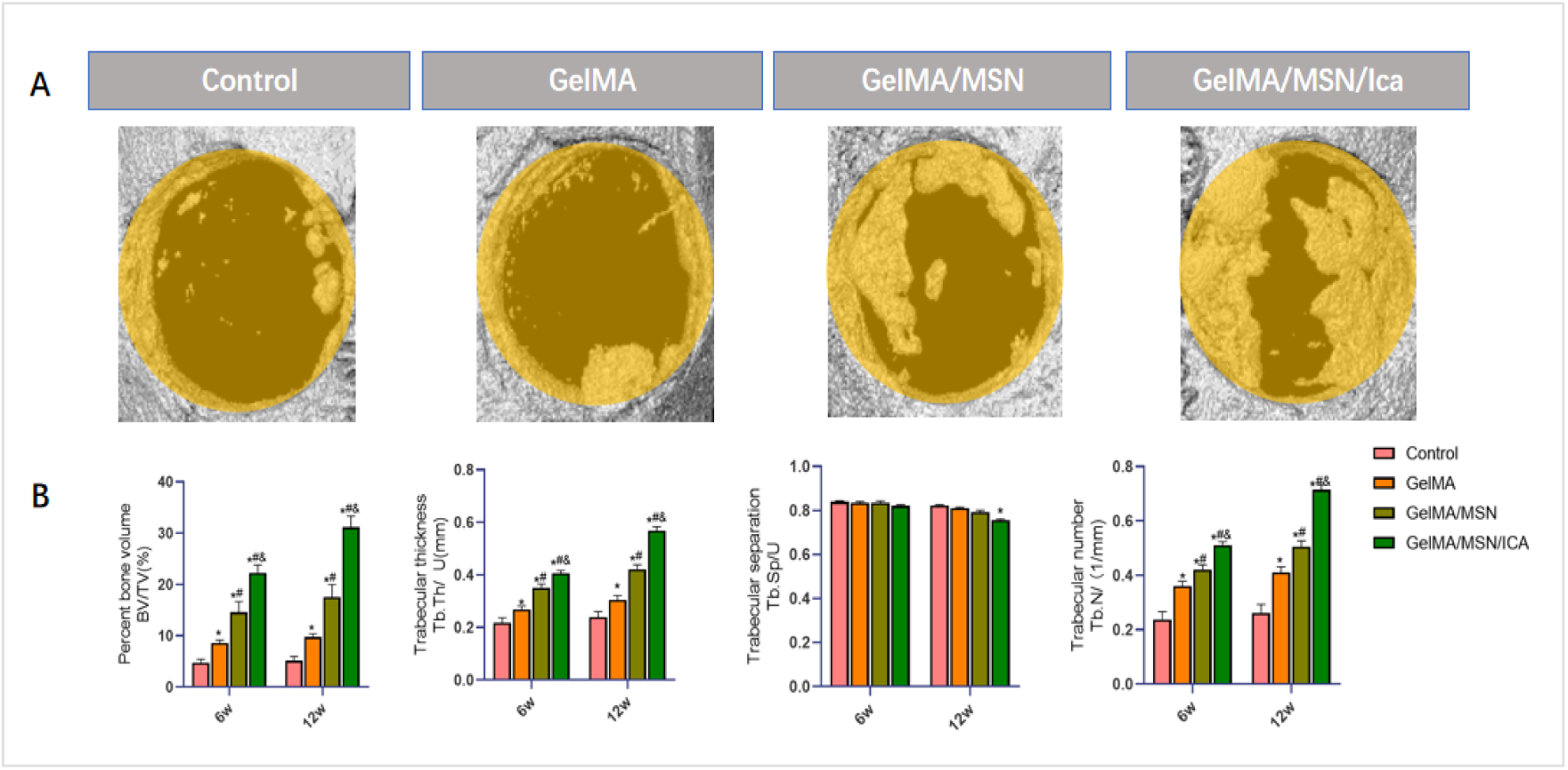
(A) Micro-CT Images of the skull bone defect at 12 Weeks. (B) Micro-CT Analysis of Bone Regeneration with the control group, GelMA, GelMA /MSN and GelMA /MSN/ICA group at 6 and 12 Weeks (n = 3). All values are expressed as mean ± SD, n = 3.

In addition, HE staining of the skull defect at 12 weeks showed that mature trabecular bone formation was observed in the GelMA/MSN/ICA group, which was more significant than that in the GelMA/MSN group, while fibrous tissue infiltration was observed in the blank group (Fig 7A). After decalcified sections of bone tissue, immunofluorescence staining was performed to directly show the local bone regeneration, and the bone regeneration marker Col1 was detected (Fig 7B). It can be seen that the signal intensity of GelMA/MSN/ICA group was significantly higher than that of other groups, and the expression of Col1 was better than that of other groups.

**Fig. 7.**
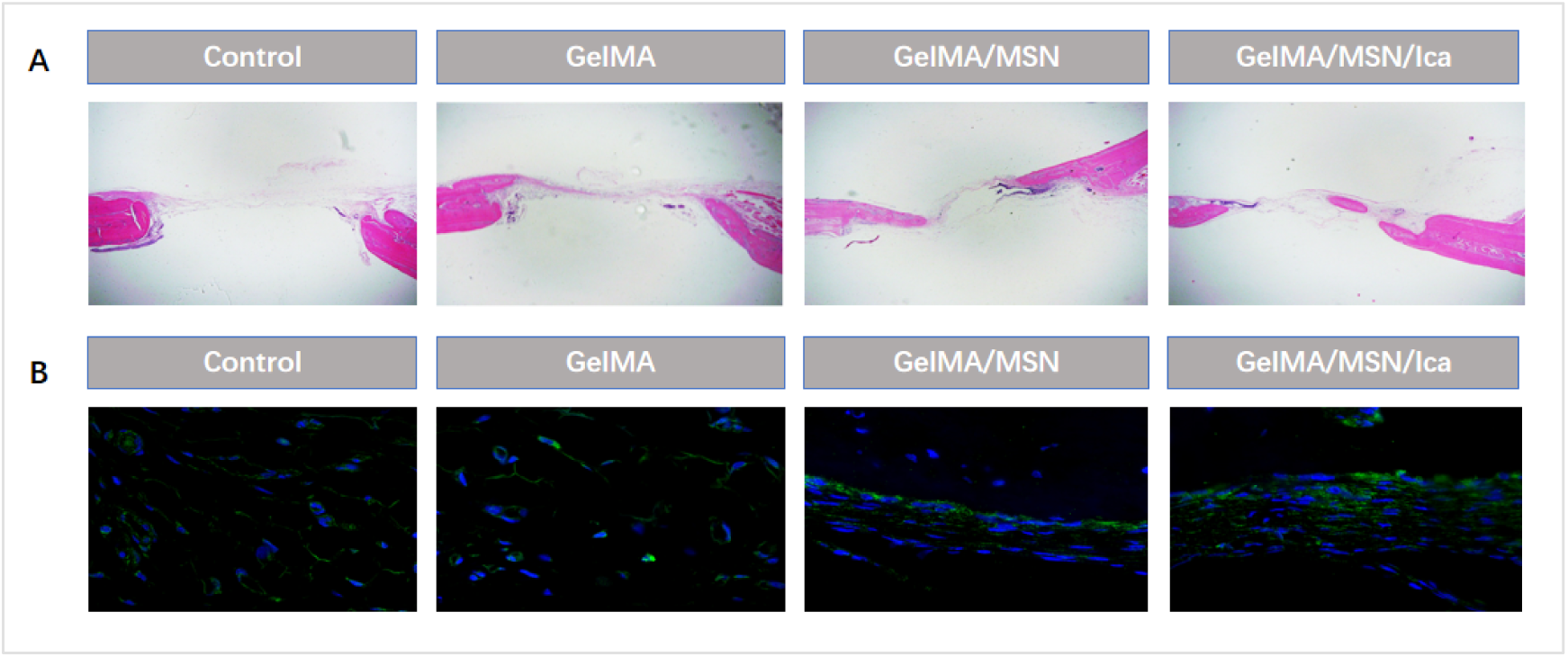
(A) HE staining of the skull defect at 12 weeks. (B) Immunofluorescence staining of Col1(Green). n = 3

## 4. Discussion

Bone defect caused by trauma, tumor resection or congenital malformation is a major clinical challenge, and millions of patients worldwide seek treatment every year. Traditional treatment methods, including autologous transplantation and allogeneic transplantation, are limited by the incidence of donor site, immune rejection and insufficient mechanical stability, while synthetic materials (such as ceramics and metals) often lack biological activity and cannot simulate the dynamic bone remodeling microenvironment[25–27]. Gelatin methacryloyl (GelMA) is a photocrosslinking hydrogel derived from extracellular matrix (ECM). Due to its adjustable physical and chemical properties, biocompatibility and ability to support osteogenic differentiation, GelMA has become a promising candidate for bone tissue engineering[28, 29]. However, GelMA based scaffolds exhibit suboptimal mechanical strength, rapid degradation and limited osteoinductive ability, which hinder their clinical conversion to critical size defects [30]. In this study, a composite hydrogel composed of MSN and GelMA with ICA was developed. The composite system has stable microenvironment for cell proliferation and differentiation, stable and controllable degradation rate, appropriate mechanical properties, and good drug release efficiency, so as to promote bone regeneration and become an effective solution for bone tissue engineering.

Our GelMA/MSN/ICA composite hydrogel system solves these limitations through rational integration MSN and ICA. Studies have shown that the density range of human cancellous bone is 75-90%, and the unit diameter is about 50-300 μ M[31]. GelMA hydrogels can also be freeze-dried to produce porous scaffolds with controllable pore size and porosity [32]. For example, Chen et al.[33] synthesized GelMA hydrogels with different degrees of substitution (49.8, 63.8 and 73.2%) using 1, 5 and 10 m Ma solutions, respectively. In these experiments, the average pore size of GelMA hydrogel obtained from freeze-dried GelMA gel characterized by SEM is 50 mm (49.8%), 30 mm (63.8%) and 25 mm (73.2%), so the pore size of the composite hydrogel matches the human bone parameters in the physiological environment, making it closer to the physiological state and providing a good space for nutrient exchange[22]. In addition, Liu et al reported that the MSN doped polycaprolactone scaffold increased the compressive strength by 40% and continued to release drugs [34].The rheological test showed that the composite hydrogel had good injectability and turned into a good gel state under UV photocrosslinking. In addition, the swelling ratio of the hydrogel showed that the composite hydrogel with MSN had good crosslinking density. EDS analysis showed that Si ions were evenly distributed on the whole composite scaffold, synergistic osteoimmunomodulation via silicic ions and ICA. Moreover the uniform distribution of MSN made the hydrogel surface have a certain rough contact surface, which enhanced the mechanical strength of the hydrogel, and the irregular pore geometry and solid structural integrity were also maintained. The regular pore results and the mixing of MSN provided rough indication and sufficient space for cell adhesion, providing the best microenvironment for cell adhesion and nutrient transport [35].

The degradation of the composite hydrogel system tended to be stable after 16 days, and the degradation was incomplete, possibly because the addition of MSN enhanced the direct adhesion between hydrogels and cells, and the residual composite hydrogel provided a local extended support structure without hindering bone remodeling. Some studies have shown that the growth factor BMP-2 loaded into MSN shows obvious osteogenic differentiation of mesenchymal stem cells [36]. This composite hydrogel carries ICA into MSN. It is found that the continuous degradation of hydrogel is conducive to the release of MSN, thus promoting the continuous release of ICA. The initial burst and continuous release of ICA can be attributed to the mesoporous structure of MSN, which physically adsorbs ICA and realizes PH/enzyme response diffusion[13, 37]. The sustained release of ICA in the later stage may be related to the slow degradation of hydrogel. This sustained-release drug meets the phased requirements of bone repair. Early inflammation (ph∼6.5) triggers MSN pore expansion to accelerate drug release, and then gradually administered in the late remodeling stage, The initial burst release (52.7% at 24 h) may suppress early inflammation[38, 39].

In previous studies, GelMA is often used as 2D or 3D cell culture matrix material due to its inherent high biocompatibility[40], and contains many RGD sequences that can promote cell adhesion[41, 42]. Studies using graphene oxide reinforced GelMA obtained similar mechanical properties, but reported cytotoxicity at high filler concentrations (>7 wt%) [43].In this study, the results of phalloidin/DAPI staining analysis showed that the rat bone marrow mesenchymal stem cells in each group had high ductility and showed bright and orderly arranged filaments. Therefore, we believe that the addition of ICA and MSN will not have a negative impact on the adhesion properties of BMSCs. In addition, CCK-8 test and live/dead staining results showed that the number of cells in the composite gel group was significantly higher than that in the other two groups by the fourth day, and the cultured cells showed vigorous growth, indicating its excellent biocompatibility. On the contrary, the proliferation rate of bone marrow mesenchymal stem cells in GelMA group was the slowest because of the lack of MSN and ICA. The decrease of the growth rate of rBMSCs in the later stage may be due to the limited space in the cell culture plate and the inhibition of cell contact. In conclusion, our findings indicate that it provides a supportive environment for cells and effectively promotes cell proliferation.

ALP and ARS staining of osteogenic differentiation showed that ALP was detected and analyzed quantitatively on the 7th day. GelMA /MSN/ICA composite hydrogel system significantly enhanced osteogenic differentiation. On the 14th day, ARS staining showed that mineralized and calcified nodules were the most obvious, which was in the GelMA group alone. Therefore, we found that the released ICA could effectively promote osteogenic transformation within 14 days. The expression levels of ALP, Runx2 and col1 in rat BMSCs were detected by RT qPCR. We found that the composite hydrogel group was the highest on the 7th and 14th day, which also showed that it effectively promoted bone differentiation. ALP is the key marker of early osteogenesis, representing the early differentiation of BMSC into osteoblasts [44]. Col1 is one of the main components of bone matrix, which can promote the adhesion and differentiation of osteoblasts. Col1 is a key osteogenic ECM component, and its upregulation highlights the ability of hydrogel to promote bone matrix synthesis and mineralization[45]Studies have shown that, in terms of mechanism, the released ICA activates ER α-Wnt/β-Catenin and Bmp-2/Smad pathways, which is consistent with the established dual role in enhancing osteoblast differentiation and inhibiting osteoclast activity[46, 47].In addition, the sustained silicic acid release from MSN degradation synergistically activates Wnt/β-catenin with ICA[48, 49]. In general, GelMA/MSN/ICA composite hydrogel can effectively promote the osteogenic differentiation of rat bone marrow mesenchymal stem cells in *vitro*.

Animal experiments showed that after skull defect modeling and hydrogel implantation, Micro-CT showed that the composite hydrogel group showed good bone regeneration at the bone defect interface at 6 and 12 weeks. HE staining showed the histomorphology of the bone defect, and the formation of mature trabecular bone was observed in GelMA /MSN/ICA group. The results of HE staining were consistent with the trend observed by Micro-CT scanning. The expression level of Col1 by immunofluorescence staining was consistent with the results of gene expression analysis at the cell test level, indicating that Col1 effectively stimulated bone regeneration at the implant interface.

Several limitations constrain current interpretations, First of all, the residual silica nanoparticles (<5 wt% at 12 weeks) will not cause acute toxicity, but long-term biodistribution research is needed, because it is reported that silica has accumulated in the liver and spleen for more than 6 months[50]. Secondly, although rat skull defects provide a standardized model, they are difficult to replicate mechanical loads in human load-bearing bones[51]. he validation of dynamic compression test in large animals (such as sheep tibia) is guaranteed. Thirdly, although MSN can make ICA release continuously, the sudden release within the first 24 hours may require surface modification for stricter time control.

## 5. Conclusion

In this study, we mixed MSN with ICA and GelMA hydrogel to form a composite hydrogel to enhance bone regeneration and bone integration. MSN made the gel more stable and functional. ICA carried on MSN was evenly distributed in the composite hydrogel and could be released for more than 15 days. The sustained release of ICA locally stimulated the proliferation and osteogenic differentiation of rBMSCs, and improved the early osseointegration. Notably, this study first reports MSN-mediated immunomodulation in GelMA hydrogels for bone repair, addressing a critical gap in natural polymer composites.

## Author Contributions

Y.Z. and C.Z. contributed to the conception and design of the study. C.G., X.Z., Y.Z., Y.T., H.L., Y. J and X.C. contributed to the data acquisition, interpretation, and analysis. Y.L., and L.F. contributed to the data interpretation and revision of the manuscript. All authors approved the submission of the paper.

## Acknowledgements

This work was supported by Henan Province 2025 Science and Technology Development Plan (Grant Number:252102310375). Special Project for Scientific Research of Traditional Chinese Medicine in Henan Province (Grant Number: 2023ZY2121).

## Conflict of interest

The authors declare that they have no known competing financial interests or personal relationships that could have appeared to influence the work reported in this paper.

## Notes

### Competing Interest Statement

The authors have declared that no competing interests exist.

